# Using Propensity Score Matching to Control for MRI Scan Quality

**DOI:** 10.1101/2025.09.26.678901

**Authors:** Veronica J. Cramm, Tyler M. Call, John A. E. Anderson

## Abstract

Movement during MRI scanning complicates distinguishing between the different tissues in the brain (e.g., grey and white matter). Standard practice excludes scans based on researcher-determined visual quality thresholds. Unfortunately, children, elderly, and clinical populations are shown to move more, resulting in higher exclusion rates. This disproportionate exclusion creates systematic bias in the literature and makes research findings less generalizable. Furthermore, the artifacts caused by motion are demonstrated to continue to confound data, even after visual quality control has occurred. We aimed to minimize the confounding factor of systematic group differences in movement. To achieve this, we used a post-scanning statistical technique called propensity score matching (PSM) that matches control and patient populations on scan quality metrics, leading to more comparable groups, greater inclusion, and more generalizable results. We found that PSM can attenuate significant differences in scan quality between groups while allowing for greater sample diversity than standard exclusion protocols. Crucially, using PSM can also alter the results of neuroimaging-based analyses. Using three datasets (total n = 1536), we compared voxel based morphometry analyses based on different quality control protocols. In particular, we observed discrepant results between PSM and strict threshold exclusion, with PSM magnifying some regional group differences and diminishing others. Overall, PSM is a customizable way to mitigate the impact of confounds in neuroimaging research and a powerful method to help distinguish true effects from artifacts.

## Introduction

Participants in MRI studies are required to remain as still as possible, and yet, small movements are still very common. Such movement degrades image quality by introducing motion artifacts such as ringing and blurring (Gilmore et al., 2021). These movement artifacts complicate inference in structural MRI studies by introducing partial voluming effects. Current best practice for quality control (QC) of structural MRI scans relies on exclusion of poor quality scans to mitigate motion bias (Gilmore et al., 2021). However, movement is not entirely random (Gilmore et al., 2021; Van Dijk et al., 2012): it is more common in men (Alexander-Bloch et al., 2016; Van Dijk et al., 2012) and clinical, pediatric, and elderly populations (Hausman et al., 2022; Pardoe et al., 2016; Satterthwaite et al., 2012). For instance, individuals with schizophrenia and Autism Spectrum Disorder (ASD) move more than age-matched healthy controls (Pardoe et al., 2016). Additionally, higher clinical severity is associated with more movement artifacts (Nakua et al., 2023). Therefore, rigorously excluding high-movement individuals from a study likely disproportionately removes the most clinically impacted individuals and introduces systematic biases. We aim to address this issue by using *propensity score matching*, a novel approach in this field, allowing researchers to include a wider range of clinically affected individuals across a more diverse set of scans to improve the generalizability and replicability of MRI studies.

In-scanner motion affects functional, structural, and diffusion MRI (Alexander-Bloch et al., 2016; Van Dijk et al., 2012; Yendiki et al., 2014). Movement of fractions of a millimeter has been shown to drive functional connectivity differences between groups, even after applying standard rigid body transformations to correct for motion (Van Dijk et al., 2012). In structural MRI, such as T1-weighted contrasts, most segmentation errors occur as a result of partial voluming effects (Smith & Nayak, 2010; Zaitsev et al., 2015), which occur when more than one type of tissue is present within a single voxel. Partial voluming effects increase when motion blurs the boundaries between tissue types. For example, Reuter et al. (2015) showed that as motion increased, measurements of gray matter volume (GMV) decreased along the gray matter and CSF boundary and increased along the gray matter and white matter boundary. To mitigate motion effects and partial voluming in functional data, rigid body transformations are standard. However, this approach cannot typically be applied to structural scans since, typically, only a single image is acquired. As an alternative, some studies have explored using proxy motion metrics (e.g, from functional scans) to regress out movement in structural scans (Van Dijk et al., 2012), but the reliability of this method is unclear.

Due to the unclear efficacy of motion regression, the standard approach is still to exclude poor quality scans. To determine scan quality, expert raters typically conduct visual inspections of MRI data on a variety of metrics including blurring or ringing. Thresholds for each metric are set by the research team on a study-by-study basis to determine which scans to include for analysis (Gilmore et al., 2021). Notably, there are currently no standard approaches for determining acceptable thresholds for noise, which may vary in several ways: by dataset, e.g., if a clinical population is extremely scarce, thresholds may be lower in order to include more individuals; research question, e.g., if the researcher is more interested in total brain volume or thickness measures; and researcher background, e.g., clinicians may tolerate higher-movement scans relative to researchers studying healthy populations.

Visual inspection and threshold exclusion is a problematic approach for assessing neuroimaging quality as it can introduce systematic bias (Nakua et al., 2023). These varying criteria within and across studies can drive systematic differences between vulnerable populations and healthy controls. Furthermore, within the context of a single study, there is a tension between the impact of noise and a high inclusion threshold which can reduce power by excluding participants and reducing sample sizes. All of these factors reduce generalizability. Furthermore, variability in findings across certain literatures featuring a high degree of MRI noise makes it hard to compare and synthesize results. While visual quality ratings are significantly correlated with automated quality control measures (*R^2^* = 0.90; Gilmore et al., 2021), there is a real concern that visual inspection may miss some indicators of poor scan quality, leading to a search for more informative and reliable automated QC methods (Baum et al., 2018; Pardoe et al., 2016). Given the lack of a consistent approach to MRI scan quality, there is a need to identify a method that adjusts bias across datasets and can be applied consistently. In this study, we attempt to find a balance between these extremes that minimizes confounds and maximizes generalizability.

We propose propensity score matching (PSM) as a solution that controls for noise while preserving the clinical features of the population being studied. PSM matches participants based on the probability of belonging to a specific group by using logistic regression to predict out-group membership. Specifically, PSM equates groups based on covariates (such as movement or scan quality) by balancing based on the propensity score (distance) between groups (Baek et al., 2015). Balancing the confounds makes the two groups more comparable, thus allowing high movement populations to be better represented in the study, as scan quality is now controlled between groups. Our study will use automated measures of scan quality from the CAT12 toolbox (e.g., resolution, noise, bias, mean surface Euler number, defect area, weighted average (IQR)) to avoid subjective rater bias and balance the groups. These metrics from CAT12 have been shown to match the performance of visual inspection (Gilmore et al., 2021). Additionally, using these automated measures removes the subjectivity of human rating, and therefore may be more consistent. Thus, we propose the use of automated QC measures and PSM to maintain diversity within datasets while mitigating motion bias. We predicted that 1) The groups (e.g., autism vs. controls, schizophrenia vs. controls, and older vs. younger adults) will have significant differences in scan quality prior to quality control; 2) these differences will drive the group differences in GMV; and finally 3) PSM will attenuate some or all of the differences between groups.

## Materials & Method

### Participant and MRI Acquisition Details

Participants from three open source datasets were separately obtained and analyzed for this study. Healthy aging, autism and schizophrenia datasets were acquired from the Neurocognitive Aging data release (Spreng et al., 2022), Autism Brain Imaging Data Exchange I (ABIDE I; Di Martino et al., 2014), and Centers of Biomedical Research Excellence (COBRE; Aine et al., 2017; Çetin et al., 2014), respectively. Data for the healthy aging and clinical groups were obtained from OpenNeuro and the Collaborative Informatics and Neuroimaging Suite (COINS; Landis et al., 2016; Scott et al., 2011; http://coins.mrn.org). All datasets consisted of a treatment group and accompanying controls (see Table 1 for participant demographic information and Figure 1 for details on participant removal). We acquired structural T1-weighted (T1w) scans for each dataset (see Table S1 in supplementary materials for MRI acquisition details). Further information can be found on the datasets’ respective websites^1^.

**Figure 1.**
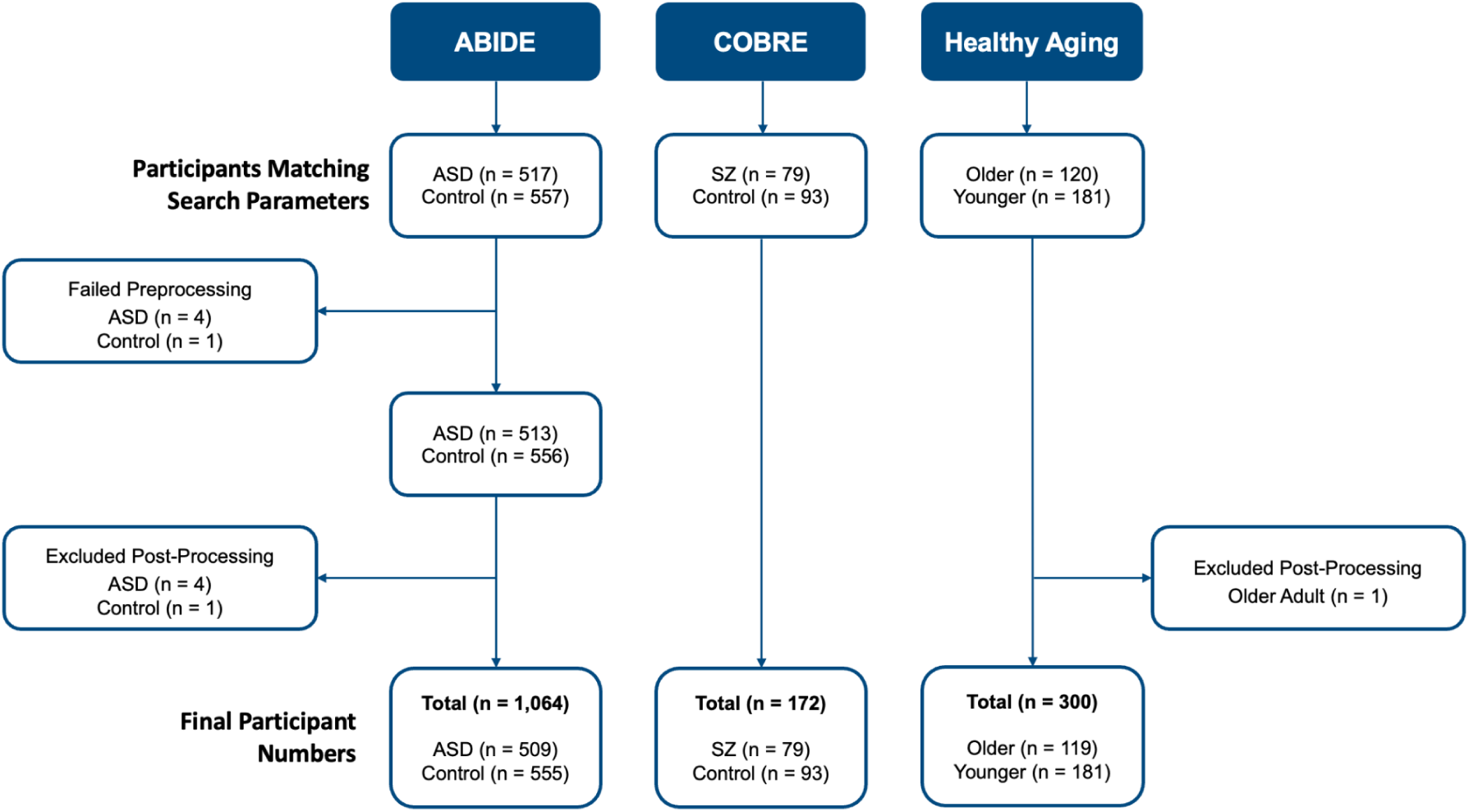
Overview of participant removal. *Note.* Post-processing removal is due to identification of poor quality scans after preprocessing concluded or scans that interfered with successful implementation of Combat harmonization software.

**Table 1.** Overview of final participant demographics by dataset and analysis level.

| **Dataset** | **Analysis** | **N (Patient / Control)** | **Age Range (Years)** | **Sex (M/ F/ O)** |
| --- | --- | --- | --- | --- |
| **ABIDE** | Full Inclusion | 509 / 555 | (6.47, 64) | 908 / 156 |
|  | Exclusion | 445 / 516 | (6.47, 58) | 818 / 143 |
|  | Matched | 497 / 497 | (6.47, 64) | 850 / 144 |
| **COBRE** | Full Inclusion | 79 / 93 | (18, 65) | 131 / 41 |
|  | Matched | 79 / 79 | (18, 65) | 131 / 37 |
| **Healthy Aging** | Full Inclusion | 119 / 181 | (18, 89) | 132 / 167 / 1 |
|  | Matched | 115 / 115 | (18, 89) | 101 / 128 / 1 |

*Note.* Sex was entered as male (M), female (F), or non-identifying (O) with standard labels.
<sup>1</sup> https://openneuro.org/datasets/ds003592/versions/1.0.13 (Healthy Aging) https://fcon_1000.projects.nitrc.org/indi/abide/ (ABIDE) https://fcon_1000.projects.nitrc.org/indi/retro/cobre.html (COBRE)

### Image Processing

Images from all three datasets were subjected to the same pipeline, including preprocessing, quality metric extraction, harmonization, and voxel based morphometry (VBM; see Figure 2 for pipeline overview). To begin, T1w images were preprocessed using the standard Computational Anatomy Toolbox (CAT12; https://neuro-jena.github.io/cat/) for Statistical Parametric Mapping (SPM) in MATLAB (Gaser et al., 2024). The preprocessing pipeline consisted of two main steps. In the initial processing step, spatial adaptive non-local means (SANLM) denoising and bias correction were applied to the raw images to reduce noise and correct for field inhomogeneities, thereby ensuring more accurate results that reflect the true brain structure for each participant. The second step involved skull-stripping, parcellating, and registering the brain to a 1.5mm^3^ voxel size MNI152 template provided by the developers. The result of this refined processing included three main segmented images: grey matter (GM), white matter, and cerebrospinal fluid. For the purposes of this study, only the grey matter segmentation was used in subsequent analyses.

**Figure 2.**
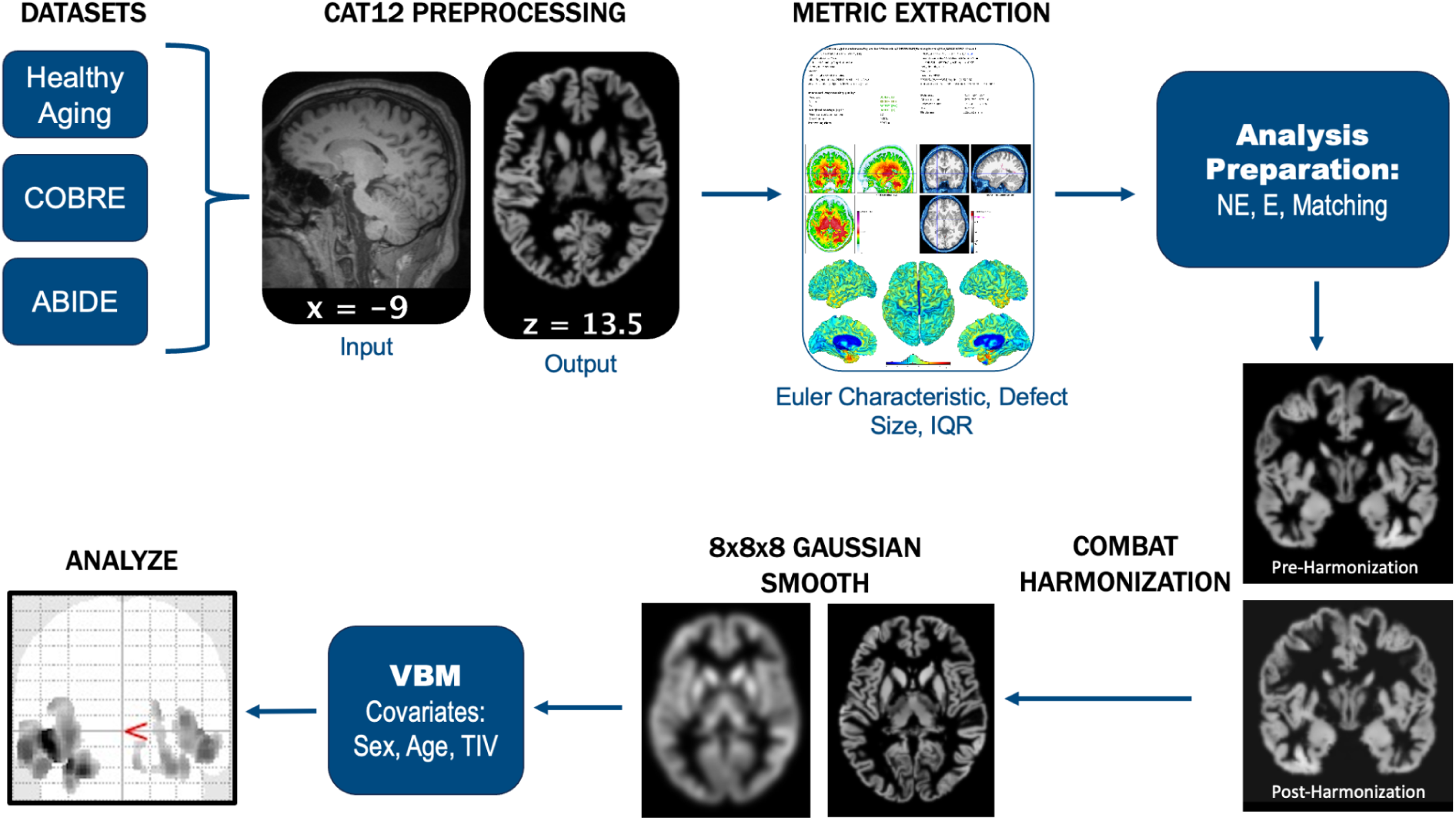
Graphical depiction of the standardized preprocessing pipeline used for all datasets.

Prior to undergoing VBM using SPM12 (https://www.fil.ion.ucl.ac.uk/spm/software/spm12/; Penny et al., 2011), modulated-warped GM images were smoothed with a standard 8mm^3^ Gaussian kernel. Smoothing helps to improve the signal-to-noise ratio (SNR) and normalize individual differences across participants (Ashburner & Friston, 2000). Healthy aging and ABIDE datasets were harmonized to account for site differences before smoothing.

### Quality Assessment

CAT12 automatically generates several quality control metrics as part of preprocessing: resolution, noise, and bias, which were averaged into an overall image quality rating (IQR). *Resolution* refers to the original acquisition resolution voxel size. Larger voxel sizes are rated lower since they miss fine detail and have a higher risk of partial voluming effects, i.e., of a single voxel containing more than one tissue type. *Noise*, also known as thermal or Nyquist noise, refers to the motion of electrons—in the MRI hardware and the participant’s body—that degrades the SNR (Aja-Fernández & Vegas-Sánchez-Ferrero, 2016). *Bias* refers to the magnitude of the bias field change across the MR image. Higher levels of bias normally cause part of the image to appear less intense (darker) than other regions (Juntu et al., 2005).

When reconstructing the surface, prior to correction, CAT12 also produces surface metrics that complement the above volume-derived metrics. These metrics include the mean surface *Euler number*, which captures the complexity of the gray/white matter boundary and is highly sensitive to noise; and *defect size*, the percentage of the brain impacted by topological defects (Gaser et al., 2024).

### Scanner Harmonization

Where studies were collected across multiple scanners, we removed the effect of scan site using the Matlab implementation of NeuroCombat (Fortin et al., 2017) to harmonize the GM images. Sex, age, and group were included as covariates to preserve natural biological variability. The segmented GM files were vectorized and combined into a matrix within Matlab (https://www.mathworks.com; R2023a). The data were harmonized and the resulting vectors were converted back into segmented GM images. These harmonized images were then smoothed in CAT12 prior to VBM.

### Quality Control Protocols

Three quality control protocols were tested to explore the full impact of quality control on standard group comparisons. First, we used a full-inclusion protocol, in which all successfully preprocessed participants were included for analysis. Second, we used an exclusion protocol, in which all participants below 70% IQR were excluded. This threshold was chosen based on the recommendation of the CAT12 documentation (Gaser et al., 2024) and is meant to represent strict exclusion. Finally, we used the PSM protocol (see below), in which control and patient groups were matched on three CAT12 quality assessment metrics; IQR, Euler characteristic, and defect percentage.

### Propensity Score Matching

Propensity score matching was completed in R (R Core Team, 2021) using the MatchIt package (Ho et al., 2025). The propensity score, representing the probability of group assignment given the observed covariates, was estimated using a logistic regression model with *Euler characteristic*, *defect size*, and *IQR* as the matching metrics.

Matching was performed using a 1:1 optimal matching algorithm, ensuring balance in covariate distributions and equal sample sizes between groups. Post-matching balance was assessed using standardized mean differences (SMDs) and visualized with Love plots to confirm that improved balance was achieved. An SMD of less than |0.1| was considered ideal. For consistency, all datasets underwent the same matching procedure, regardless of whether ideal balance was achieved.

### Voxel-Based Morphometry (VBM)

GM images were analyzed in SPM12 using general linear models (GLMs) to compare groups while controlling for relevant covariates (total intracranial volume (TIV), sex, and age where applicable). Three analyses were conducted for each dataset enabling us to compare whether PSM was more effective at detecting group differences and reducing the impact of noise than standard exclusion or no correction.

We used the cluster-correction method from SPM12 (Gaser et al., 2024; Penny et al., 2011), which uses a two-step cluster-correction procedure that starts with a voxel-level correction (FWE, *p* < .05), followed by retention of any clusters with peak voxels that were distinct at 8mm apart.

### Between-Method Comparisons

Two-sample t-tests were conducted using base R, for each QC protocol, to determine if there was a significant difference in image quality prior to PSM, and furthermore, if either exclusion or matching protocols attenuated this difference.

## Results

### Propensity Score Matching

Propensity score matching improved balance across all metrics for all datasets to varying degrees (see Figure 3). Matching was most successful for the ABIDE dataset, achieving an SMD below the |0.1| threshold for all metrics, indicating strong balance between groups. Additionally, PSM allowed for the inclusion of 52 more participants with autism compared to traditional exclusion methods (Table 1), which allowed for greater inclusion of lower quality scans from both patient and control groups (Table 2). Unlike the ABIDE dataset, matching was demonstrated to be more complicated for the COBRE and healthy aging datasets.

**Figure 3.**
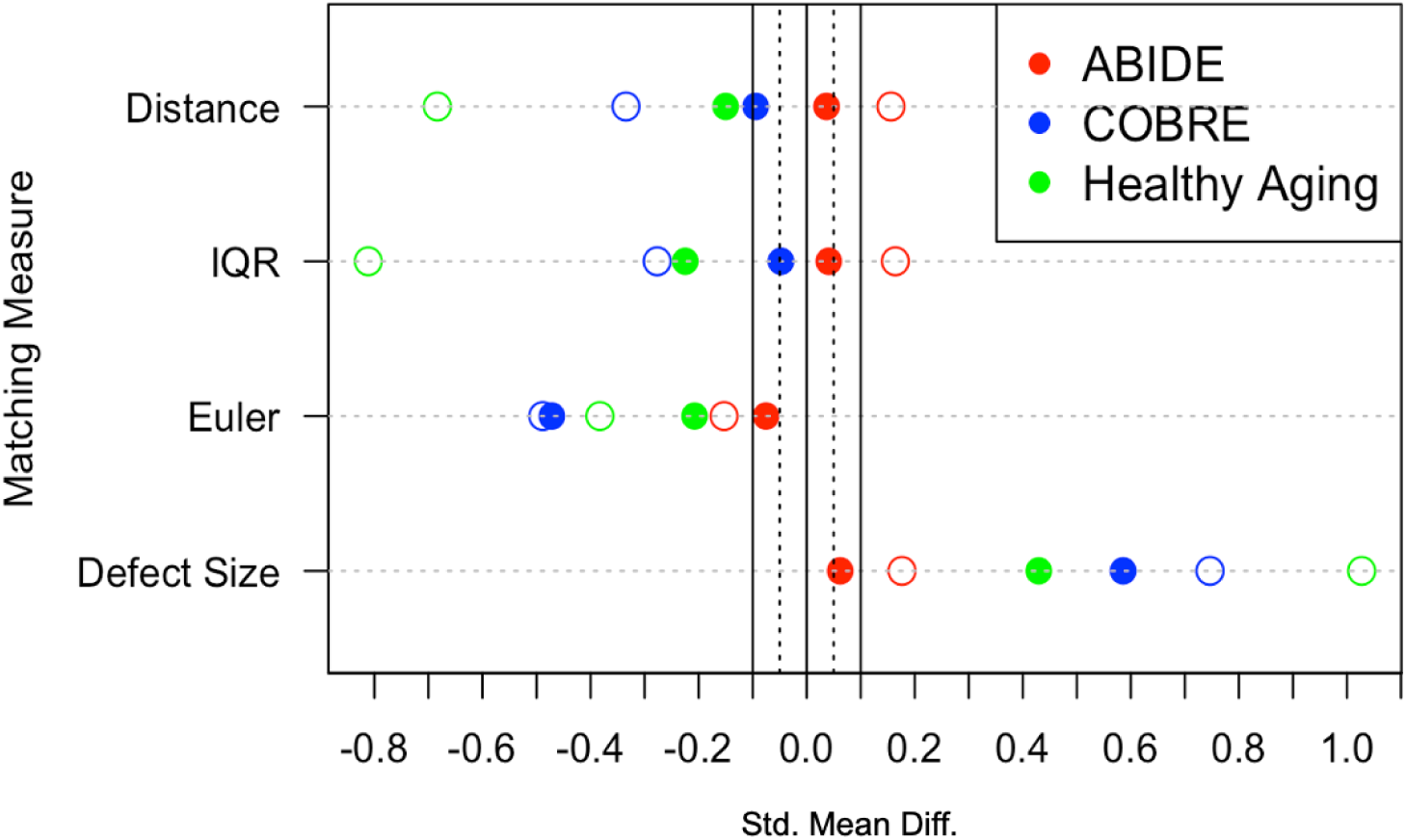
Love Plot Demonstrating Improved Covariate Balance After Matching. *Note.* Standardized mean differences before and after matching for all three datasets, assessing matching quality. The solid vertical line at 0.1 indicates a successful match. The four matching measures were the *propensity score* (*distance*), the overall *image quality rating* (*IQR*), *Euler characteristic* (indicates information about the number of holes (defects) in the image), and *defect size* (%, indicates the percentage of the brain impacted by topological defects).

**Table 2.** Overview of scan quality difference by dataset and analysis.

| Dataset | Analysis | Patient IQR (%)<br>(Min, Max) | Control IQR (%)<br>(Min, Max) | t-stat | Pearson's r | p-value |
| --- | --- | --- | --- | --- | --- | --- |
| **ABIDE** | Full Inclusion | (51.07, 88.35) | (57.36, 87.72) | -2.77 | -0.085 | < 0.01 |
|  | Exclusion | (70.17, 88.35) | (70.33, 87.72) | 0.22 | 0.007 | .82 |
|  | Matched | (56.94, 86.79) | (57.36, 87.62) | -1.38 | -0.044 | .17 |
| **COBRE** | Full Inclusion | (74.45, 85.7) | (74.05, 86.27) | -3.65 | -0.27 | < 0.001 |
|  | Matched | (74.45, 85.7) | (76.47, 85.82) | -3.44 | -0.266 | < 0.001 |
| **Healthy Aging** | Full Inclusion | (76.96, 85.92) | (76.57, 86.28) | 3.56 | -0.213 | < 0.001 |
|  | Matched | (77.12, 85.92) | (76.57, 86.01) | 1.84 | -0.121 | .07 |

*Note:* Two sample t-tests were conducted in R to determine if there is a significant difference in overall scan quality between groups. All scans were above 70% image quality rating (IQR) cutoff in the COBRE and Healthy Aging datasets, and therefore there was not a separate exclusion protocol.

For the COBRE dataset, all schizophrenia patients were successfully matched to controls, but only two of four metrics were optimally balanced. Matching was effective at mitigating group differences in IQR and Distance, but could not fully attenuate the group difference in the Euler and defect size metrics.

Matching for the healthy aging dataset was the most challenging given the choice of matching algorithm. Four older adult participants had scan quality measures that were too extreme to find an ideal match, and were therefore discarded. Furthermore, while balance meaningfully improved across metrics, none of the four metrics were balanced below the SMD |0.1| threshold.

### Between-Method Comparisons

To assess the impact of different quality control protocols, we first examined whether matching reduced scan quality differences (IQR) between groups (see Table 2). Using t-tests, all three datasets exhibited significant between-group differences in scan quality *before* matching. After matching, results indicated that scan quality differences were no longer significant for the healthy aging and ABIDE datasets. For the COBRE dataset, the magnitude of the between-group IQR difference was markedly reduced, as reflected by a lower *t*-statistic, suggesting that matching mitigated the scan quality imbalance. However, a significant group difference in scan quality remained.

### Voxel-Based Morphometry

Beyond reducing overall differences in quality indices from CAT12, we were also interested in examining whether matching would also attenuate voxel-wise differences between groups.

In ABIDE, three separate VBM analyses were run for the different QC protocols (full inclusion, exclusion, matching). FWE and cluster size corrected results for the top clusters are shown in Table 3, indicating the top regions where patients with autism had greater GMV than control participants (see Figure 3 for cluster visualization). We also ran the opposite comparison to test whether controls had higher regional GMV than patients with autism, but no regions survived correction.

**Table 3.** Voxel Based Morphometry Cluster Results for the ABIDE Autism Data Across Quality Control Protocols.

| Protocol | Hem. | Region | X | Y | Z | Size<br>(Voxels) | t-stat | Cohen's d | p<sub>uncorr</sub> | p<sub>corr</sub> |
| --- | --- | --- | --- | --- | --- | --- | --- | --- | --- | --- |
| Full Inclusion | L | Sup. Temp. Gyr. | -50 | -6 | -9 | 3106 | 7.20 | 0.44 | < .001 | < .001 |
|  | L | Parahippocampus | -27 | -18 | -26 | 1032 | 6.56 | 0.40 | < .001 | < .001 |
|  | R | Sup. Temp. Gyr. | 50 | -4 | -9 | 1161 | 5.91 | 0.36 | < .001 | < .001 |
|  | R | Fusiform Gyr. | 50 | -68 | -15 | 247 | 5.46 | 0.34 | .051 | .003 |
|  | R | Parahippocampus | 26 | -24 | -20 | 671 | 5.21 | 0.32 | .003 | < .001 |
|  | R | Med. Frontal Gyr. | 6 | 16 | -22 | 182 | 5.01 | 0.31 | .088 | .004 |
|  | R | Precuneus | 3 | -63 | 27 | 127 | 4.75 | 0.29 | .148 | .008 |
| Exclusion | L | Sup. Temp. Gyr. | -51 | -9 | -4 | 3916 | 7.19 | 0.47 | < .001 | < .001 |
|  | L | Parahippocampus | -27 | -18 | -26 | 1348 | 6.40 | 0.41 | < .001 | < .001 |
|  | R | Parahippocampus | 26 | -22 | -22 | 818 | 6.22 | 0.40 | .001 | < .001 |
|  | R | Sup. Temp. Gyr. | 51 | -4 | -9 | 1379 | 5.91 | 0.38 | < .001 | < .001 |
|  | R | Fusiform Gyr. | 50 | -66 | -16 | 347 | 5.49 | 0.36 | .018 | .001 |
|  | R | Caudate | 9 | 21 | -10 | 377 | 5.25 | 0.34 | .014 | .001 |
|  | L | Caudate | -8 | 18 | -10 | 69 | 4.88 | 0.32 | .255 | .013 |
| Matching | L | Sup. Temp. Gyr. | -48 | -6 | -10 | 3631 | 7.13 | 0.45 | < .001 | < .001 |
|  | L | Parahippocampus | -27 | -18 | -26 | 1188 | 6.56 | 0.42 | < .001 | < .001 |
|  | R | Fusiform Gyr. | 50 | -68 | -15 | 373 | 5.86 | 0.37 | .025 | .001 |
|  | R | Sup. Temp. Gyr. | 50 | -4 | -9 | 1423 | 5.83 | 0.37 | < .001 | < .001 |
|  | R | Parahippocampus | 28 | -20 | -24 | 861 | 5.43 | 0.34 | .002 | < .001 |
|  | R | Med. Frontal Gyr. | 6 | 16 | -22 | 106 | 4.95 | 0.31 | .204 | .01 |

*Note:* Only regions where participants with autism had greater GMV than controls were included. Results indicate peak MNI coordinates, cluster size at 1.5mm isotropic voxels, t-statistics, Cohen's d, and both uncorrected and FWE-corrected P values. Extent thresholds (k) are empirically determined by the SPM software (kFull Inclusion = 64, kExclusion = 57, kMatching = 70). Hem = hemisphere, Sup = superior, Temp = temporal, Gyr = gyrus, Med = medial.

Overall, five brain regions (bilateral superior temporal gyrus, bilateral parahippocampus, and right fusiform gyrus) were consistently identified across all three protocols, suggesting robust group differences that were independent of protocol. Despite these regional consistencies, the effect magnitude depended on the QC method: overall t-statistic values slightly decreased from full inclusion to exclusion, then decreased further for the matching protocol, suggesting that the balance achieved through PSM reduced the magnitude of difference between control and ASD participants.

In the full inclusion and PSM protocols, individuals with ASD were shown to have greater GMV in the right medial frontal gyrus. In contrast, this difference was not visible in the exclusion protocol, suggesting a real difference was obscured by the latter protocol. Additionally, the left and right caudate were shown to have significantly greater GMV in ASD participants, as compared to controls, when the strict exclusion protocol was used.

**Figure 3.**
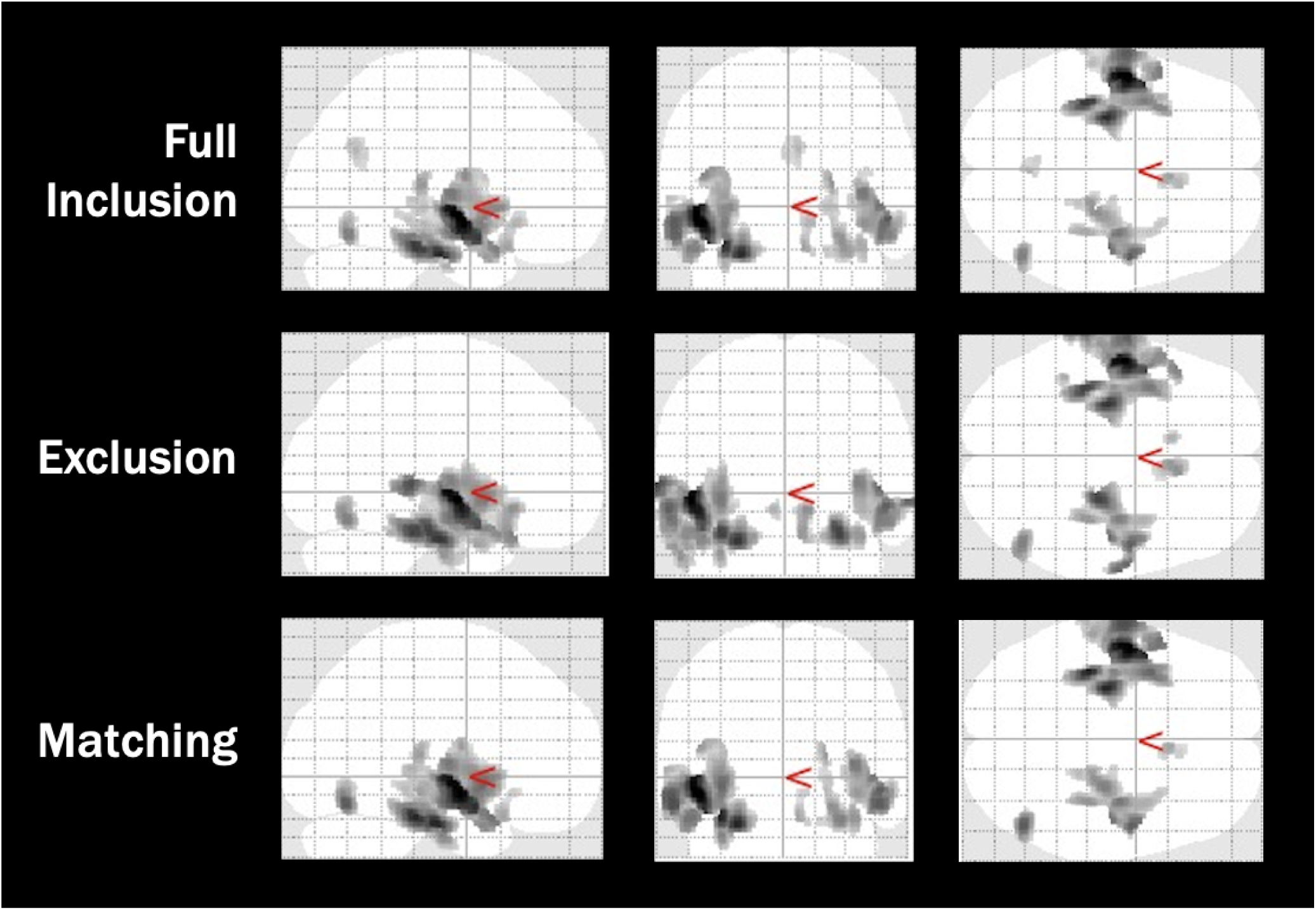
Voxel Based Morphometry Cluster Results for the ABIDE Autism Data Across Quality Control Protocols. *Note:* Images are adapted from the SPM voxel based morphometry output, indicating significant clusters where participants with autism had greater grey matter volume than controls. Output images have a resolution of 1.5mm isotropic voxels.

For COBRE, two whole brain VBMs were conducted with the full inclusion and matching protocols (see Table 4). For the COBRE dataset, there was no exclusion protocol, as all scans were above our 70% IQR cutoff. Results indicate regions where controls had greater GMV than participants with schizophrenia (SCZ). Similarly to the ABIDE dataset, the opposite analysis (SCZ > Controls) revealed no significant regions that survived correction.

**Table 4.** Voxel Based Morphometry Cluster Results for the COBRE Schizophrenia Data Across Quality Control Protocols.

| Protocol | Hem. | Region | X | Y | Z | Size<br>(Voxels) | t-stat | Cohen's d | p<sub>uncorr</sub> | p<sub>corr</sub> |
| --- | --- | --- | --- | --- | --- | --- | --- | --- | --- | --- |
| Full Inclusion | R | Inf. Frontal Gyr. | 42 | 20 | -16 | 1037 | 6.95 | 1.06 | < .001 | < .001 |
|  | L | Parahippocampus | -28 | 6 | -22 | 742 | 6.19 | 0.95 | < .001 | < .001 |
|  | R | Parahippocampus | 38 | -22 | -20 | 53 | 6.11 | 0.93 | .181 | .009 |
|  | R | Mid. Temp. Gyr. | -56 | -15 | -16 | 74 | 5.88 | 0.90 | .118 | .006 |
|  | R | Med. Frontal Gyr. | 8 | 48 | 14 | 190 | 5.47 | 0.84 | .018 | .001 |
|  | R | Inf. Frontal Gyr. | 36 | 39 | -6 | 45 | 5.26 | 0.80 | .216 | .011 |
|  | R | Mid. Frontal Gyr. | 45 | 36 | 8 | 46 | 5.06 | 0.77 | .211 | .011 |
|  | R | Uncus | 30 | -4 | -42 | 38 | 5.02 | 0.77 | .254 | .013 |
| Matching | R | Inf. Frontal Gyr. | 44 | 20 | -15 | 602 | 6.54 | 1.04 | < .001 | < .001 |
|  | L | Inf. Frontal Gyr. | -30 | 8 | -21 | 100 | 5.49 | 0.87 | .071 | .004 |
|  | L | Mid. Temp. Gyr. | -56 | -14 | -16 | 31 | 5.39 | 0.86 | .298 | .015 |

*Note:* Only regions for regions where controls had greater GMV than schizophrenia participants were included.
Results indicate peak MNI coordinates, cluster size at 1.5mm isotropic voxels, t-statistics, Cohen's d, and both uncorrected and FWE-corrected P values. All scans were above our strict exclusion cutoff (IQR<70%). As a result there is no separate strict exclusion protocol. Full inclusion and Exclusion represent the same participants. Extent thresholds (k) are empirically determined by the SPM software (k Full Inclusion = 32, k Matching = 31). Hem = hemisphere, Inf = inferior, Gyr = gyrus, Mid = middle, Temp = temporal, Med = medial.

One region, the right inferior frontal gyrus (IFG), was identified across both QC methods, suggesting the effects in this region were robust to movement. However, there was a large difference between the number of clusters identified in the full inclusion and matching protocols. Eight clusters were identified for the full inclusion protocol, whereas there were only three for the matching protocol. PSM removed regions that were less reliable/stable and overall reduced the magnitude of difference, as demonstrated by the reduction in t-statistics. Specifically, while full inclusion identified multiple clusters within the right IFG, matching retained only the most stable cluster. Additionally, matching appeared to identify clusters in the contralateral hemisphere, compared to the full inclusion protocol. While the right IFG and right middle temporal gyrus (MTG) were significantly different between groups in the full inclusion protocol, these differences were no longer significant in the matching protocol. Furthermore, the left IFG and left MTG were identified as significant in the matching protocol. In contrast, a number of regions, including the bilateral parahippocampus, right middle frontal gyrus, right uncus, and right medial frontal gyrus were only found to be significant in the full inclusion protocol.

**Figure 4.**
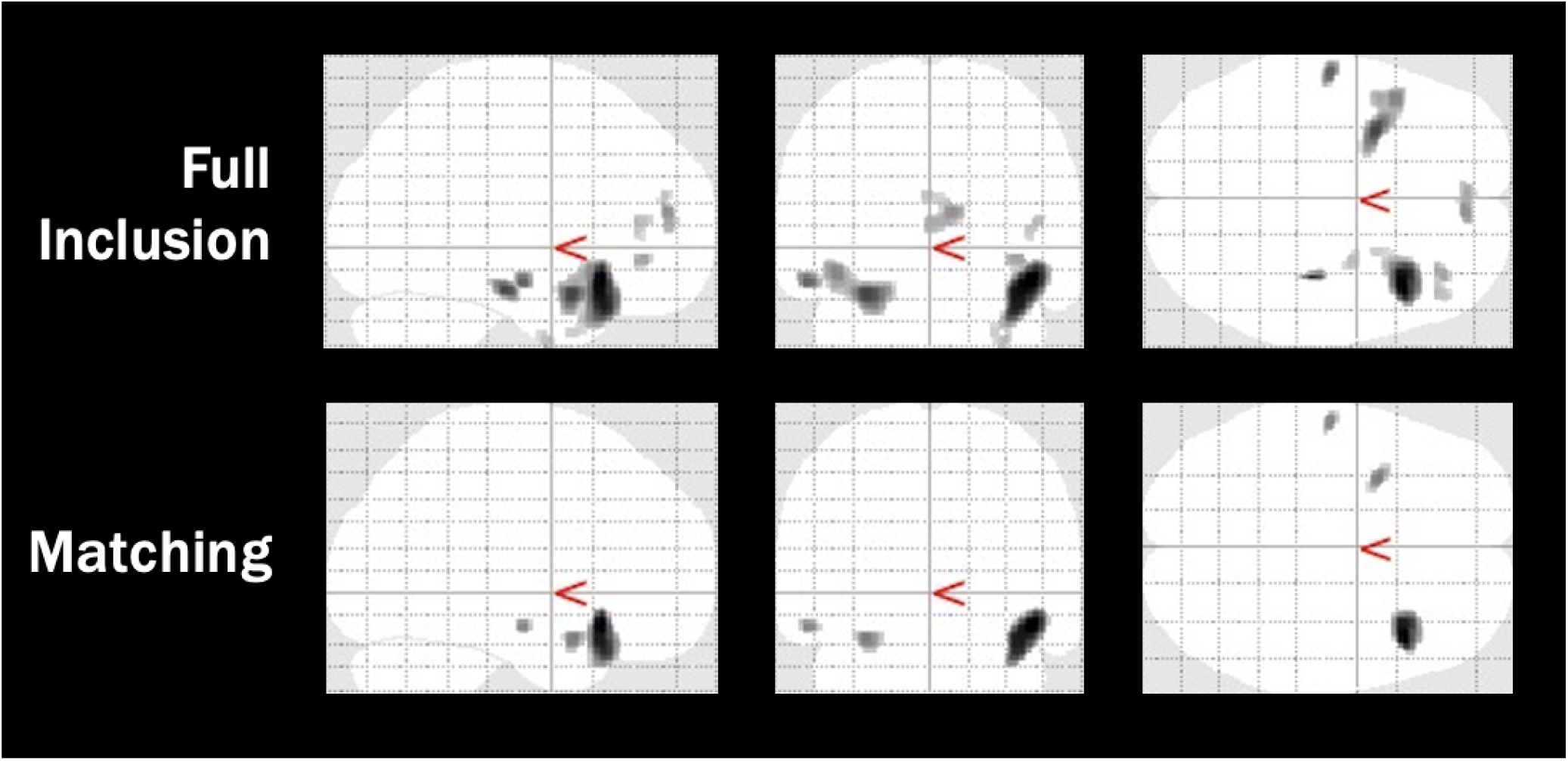
Voxel Based Morphometry Cluster Results for the COBRE Schizophrenia Data Across Quality Control Protocols. *Note:* Images are adapted from the SPM voxel based morphometry output, indicating significant clusters where control participants had greater grey matter volume than schizophrenic participants. Output images have a resolution of 1.5mm isotropic voxels.

In Healthy Aging, as with the COBRE dataset, two VBMs were conducted using the full inclusion and matching QC protocols (see Table 5 and Figure 5). For this dataset, both contrasts were found to have significant full inclusion differences. Global differences were found where younger adults had greater GMV than older adults, with the peak difference in the left anterior cingulate gyrus. Using PSM resulted in 5.34% fewer significant voxels (*n* = 22,068) than the full inclusion protocol and a decrease in the magnitude of difference at the peak of the cluster (*t*_Matched_ = 18.58, *t_Full inclusion_* = 21.63). Furthermore, across both analyses, compared to younger adults, older adults were found to have greater GMV in the left and right thalamus.

**Figure 5.**
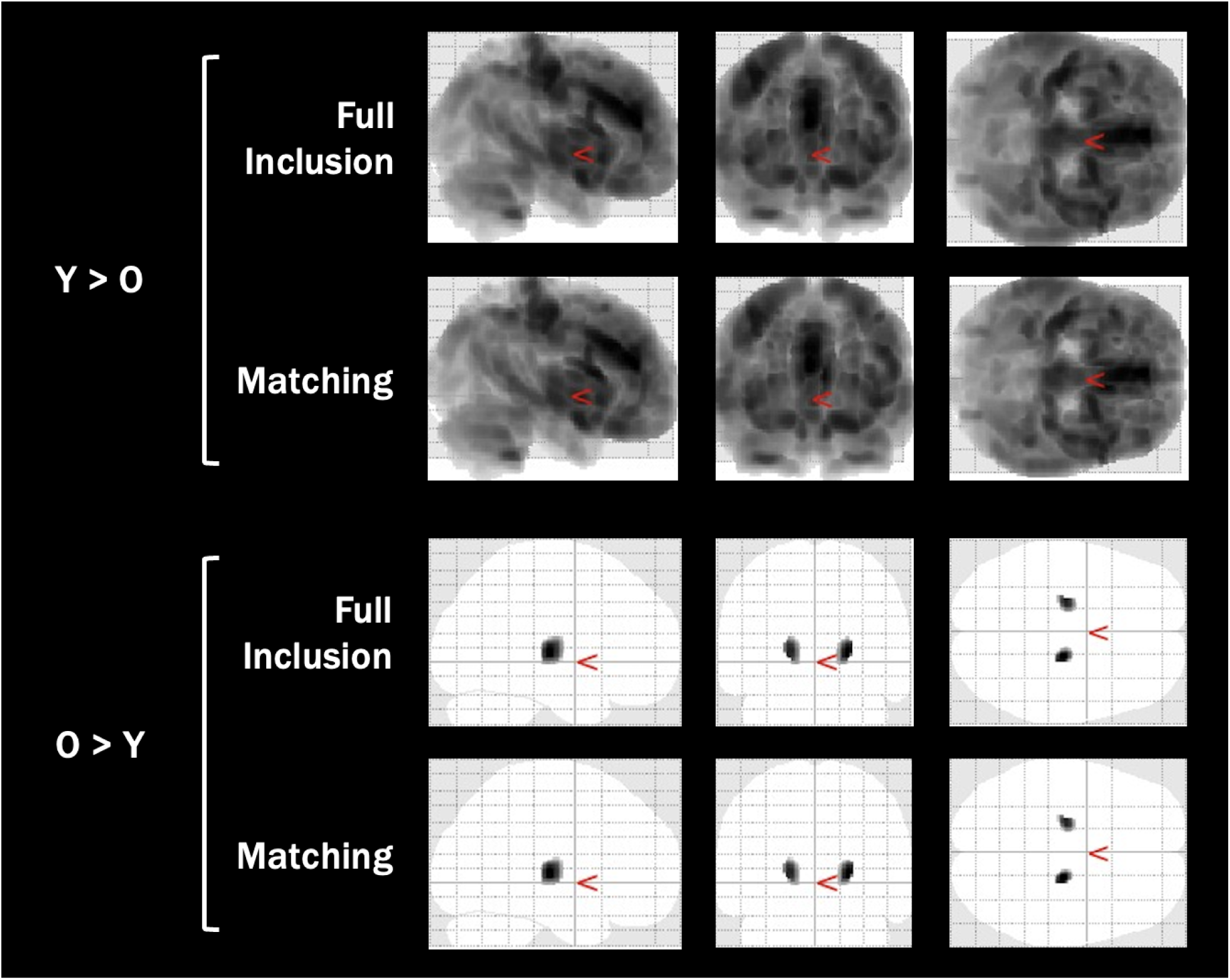
Voxel Based Morphometry Cluster Results for the Healthy Aging Data Across Quality Control Protocols. *Note:* Images are adapted from the SPM voxel based morphometry output, indicating significant clusters for both grey matter volume contrasts (younger > older, older > younger). Output images have a resolution of 1.5mm isotropic voxels.

**Table 5.**
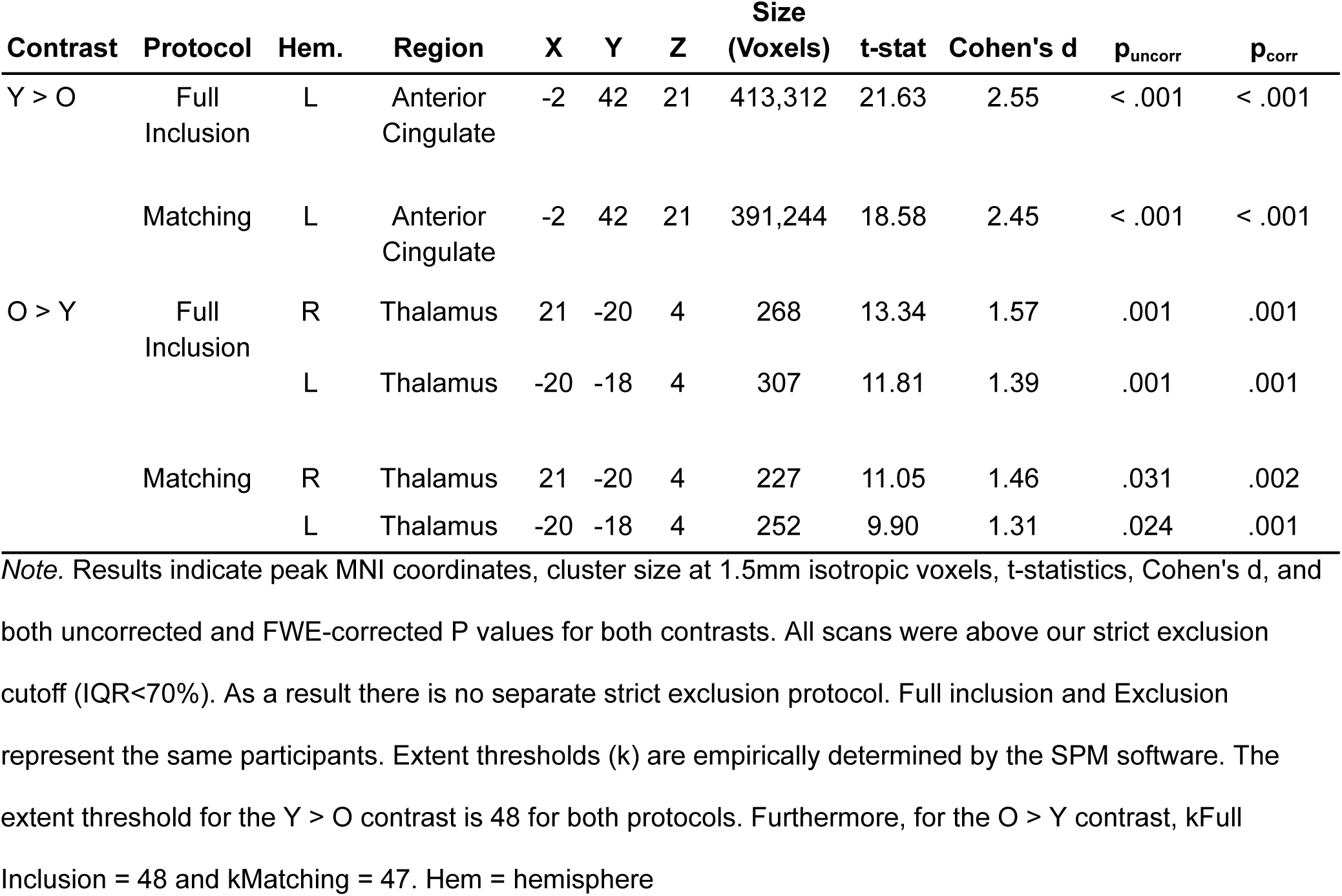
Voxel Based Morphometry Cluster Results for the Healthy Aging Data Across Quality Control Protocols.

## Discussion

This paper examined GMV and scan quality differences between control and patient groups across three datasets (healthy aging, schizophrenia, and autism) using VBM and CAT12 QC outputs. We applied typically used (threshold) quality control techniques and propensity score matching (PSM) to assess the impact of scan quality on GMV differences. We found evidence to support our hypotheses that there is a significant group difference in scan quality, prior to matching; that some of the differences in GMV are driven by scan quality; and finally, that PSM attenuates the differences in scan quality in two of three datasets and reduces the magnitude of the observed effect without sacrificing statistical power.

We sought to improve upon two techniques, visual quality exclusion and motion regression, that have demonstrated limitations. Excluding scans based on subjective ratings does not eliminate potential confounds caused by motion (Baum et al., 2018; Pardoe et al., 2016). Recently, researchers have turned to proxy regression, wherein a measure of motion collected from fMRI or dMRI is regressed out of structural data (Van Dijk et al., 2012). While growing in popularity, this method has several limitations. First, 4D images are not available in all cases and their acquisition may not always be feasible. Second, linear regression, by definition, attempts to predict the impact of a set of covariates on the data, and subsequently remove it, leaving researchers to work with residuals rather than full datasets. However, if this method’s assumption of a linear relationship between the regressed and dependent variables is not met, the residuals may not accurately reflect the unaltered data (Elman et al., 2022). Finally, because motion regression assumes that motion will be consistent across scans within a participant, it runs the risk of overcorrecting for subjects who moved more during their 4D scan than during their structural scans.

We used PSM to mitigate the impact of between-group scan quality differences while avoiding the limitations of the regression approach. Rather than regressing out the variance associated with a covariate, PSM balances covariates across groups, ultimately leading to greater cross-group comparability and unmasking differences that are otherwise suppressed with rigorous exclusion. In practice, the strength of the conclusions drawn from analyses that use PSM will depend on the nature of the balance achieved by PSM. If one manages to balance all covariates (i.e., achieve an SMD of less than |0.1| for all covariates), one can rule out these covariates as potential confounds, thus giving more weight to causal claims about the variables of interest. In contrast, if one achieves overall balance (i.e., distance or propensity score does not significantly differ between groups) but some covariates remain unbalanced, one cannot rule out that these covariates are systematic confounds, and thus causal claims are less warranted. This variability in the efficacy of PSM can be seen across our three datasets, all of which benefited from PSM but only one of which achieved ideal matching in all covariates.

Within the ABIDE dataset, PSM successfully matched all covariates within an SMD of |0.1|, allowing for greater clinical diversity to be included and subsequently highlighting contralateral regions associated with this diversity. For example, we found GMV differences (ASD > controls) in the right medial frontal gyrus (MFG) in the full inclusion and matching conditions, but not in the strict exclusion condition. In the strict exclusion analysis, both groups had higher average scan quality than the groups included in the other analyses. Assuming, consistent with findings from Nebel et al. (2022), that some low quality scans in the ASD group belong to individuals with higher clinical severity, this finding could indicate that strict exclusion removes the severely clinically impacted subset of ASD participants who influence the group differences observed in the full inclusion and matching protocols. Our results are consistent with a number of studies that also found increased volume within the right MFG for ASD participants (Bai et al., 2023; Rojas et al., 2006). However, others have observed decreased GMV in the MFG in ASD compared to controls (Hyde et al., 2009; Zhao et al., 2022).

Given that we did not find a significant difference in GMV for the left MFG, it is possible that the differences observed in other studies are inflated by systematic scan quality differences between clinical and control groups. Furthermore, Rojas et al. (2006) found that left MFG volume was negatively correlated with social impairment. While we were unable to find evidence of a similar correlation in the right MFG, we suspect that a similar correlation would obtain if impairment were analyzed. Overall, the heterogeneity in findings suggests that this region could be impacted by clinical diversity and strict exclusion protocols may systematically suppress effects within this region.

Furthermore, we observed a significant difference in GMV of the left and right caudate in ASD participants in the strict exclusion analysis. Given that caudate volume has a positive relationship with age during adolescence (Lin et al., 2015), this finding is likely the result of the increased mean age of the ASD participants in the exclusion protocol (*M* = 17.45), compared to the matching protocol (*M* = 16.89). Overall, ASD appears to be heavily influenced by clinical severity, and subsequently, scan quality, and implementing a technique like PSM can help address these confounds.

PSM appeared to have the largest impact on the COBRE dataset, in which we compared regions where controls had greater GMV than patients with schizophrenia. PSM reduces the overall number of clusters and allows for the emergence of in the contralateral hemisphere for the matching protocol. For instance, two clusters in the right IFG were identified in the full inclusion analysis, but PSM only yielded one significant cluster in that region. Additionally, PSM entirely eliminated several significant clusters from the full inclusion analysis. This elimination was particularly interesting, as all patients with schizophrenia were included in both quality control protocols, with only the number of controls varying. This finding illustrates the main benefit of PSM: by making the control and patient groups more similar, PSM included fewer control participants than the full inclusion protocol. In attempting to balance the groups, PSM trimmed the high-IQR tail of the scan quality distribution in the control group, thereby reducing the influence of having an imbalance in scan quality between groups and eliminating the inflation caused by this imbalance. Thus, PSM effectively addresses the common “apples vs. oranges” concern (Gilmore et al., 2021), in which scans that are fundamentally different in quality are compared. We demonstrated that when scan quality is considered between groups, study conclusions are dramatically altered due to a reduction in systematic group differences.

Finally, the most notable impact of PSM on the healthy aging dataset is the reduced number of significant voxels identified, with lower t-statistics at the peak. Matching was demonstrated to reduce the number of significant voxels by 5.34%, despite not achieving an ideal match. This indicates that matching attenuated some of the differences in GMV between groups, and furthermore that systematic differences in scan quality may inflate age effects. By reducing the differences in image quality, PSM reduced the variance associated with this confounding factor.

Taken together, these findings highlight the potential of PSM in neuroimaging. The flexibility and ease of implementation makes PSM an ideal quality control method to use in a field where lack of inclusivity and generalizability is a major issue. However, one caveat of PSM is that, depending on features of the dataset (e.g., distribution of covariates) to which PSM is applied, an optimal match cannot always be achieved. As we demonstrated in both the COBRE and Healthy Aging datasets, a perfect match is not always achieved. While an SMD of less than |0.1| is typically considered standard, this threshold is a guideline used for causal inference studies, and may be an unnecessarily high bar for studies that are purely aiming for improved balance.

When it is impossible to achieve ideal balance on all (or any) metrics, it is important to consider the extent of balance achieved for each metric. Less-than-ideal balance can still constitute a meaningful improvement over the unmatched dataset. If ideal balance is achieved for propensity score (distance), it indicates that *overall* the two groups are balanced. In contrast, balancing on a selected confounding variable indicates that *this* covariate is no longer a confound. In the COBRE dataset, we achieved ideal overall balance but did not balance all covariates. Despite ideal overall balance, we did not fully eliminate group differences in scan quality, as shown by t-test (see Table 2). In contrast, we did not achieve ideal balance for the propensity score or any covariates in the Healthy Aging dataset, but we still managed to reduce group differences in scan quality to a non-significant level. Thus, ideal balance does not always translate to practical improvement, and practical improvement can still be achieved in the absence of ideal balance.

One common criticism of PSM is that it reduces statistical power by removing participants. While this is a valid concern, we demonstrated in our ideally-matched dataset (ABIDE) that PSM can also increase sample size compared to strict exclusion protocols. Thus, PSM need not always result in a loss of power. Strict matching criteria may result in enough participant removal to reduce power, but more relaxed matching criteria may preserve power. Overall, these concerns should be addressed on a case-by-case basis and in the context of the research questions and samples being studied.

In this study, we chose to use the same 1:1 optimal matching algorithm for all three datasets, but researchers should choose the algorithm best suited for their data to minimize the effects of nuisance variables on their outcomes. PSM is not a one-size-fits-all technique. Depending on the research question, number of confounding variables, and tolerance for imbalance, researchers may need to try alternative matching algorithms to ours. For instance, when you are matching on several metrics and an ideal match is not achieved, we suggest implementing a method such as tiered matching, in which you match on the most important metrics first, then implement a second PSM iteration using different metrics on the intermediate matched sample. While this may result in a larger loss of participants, it would potentially allow for a smaller SMD, and therefore for causal claims to be made. In contrast, if tiered matching consistently yields sub-optimal results, adding a caliper, which reduces the allowed variance within a match, often fixes this issue (Austin, 2011; Wang et al., 2013).

Lastly, combining nearest and exact matching is another promising solution for difficult matches. Exact matching looks for pairs that are the same, which is extremely difficult with continuous variables. To combat this, you can select nearest neighbour matching, but add an “exact” parameter, where you exact match on all categorical confounders, and nearest-neighbour match on all continuous ones (Ho et al., 2025). These suggestions are some ways to handle complicated matches, but there are many more. PSM is extremely versatile and can be customized to the data.

Overall, our study was the first to apply PSM as a technique to mitigate scan quality bias in neuroimaging data. Despite this, there are a number of limitations. Firstly, we used the same matching algorithm across datasets. This choice may have compromised the efficacy of PSM for some or all of the datasets. Second, despite the large age range of participants across datasets, all scans were registered to the MNI152 brain template. Given that older adult and pediatric brains are slightly different, future work should aim to use openly available age specific templates or create a study-specific mask. Finally, our study only matched on measures of scan quality obtained through CAT12. In the future, researchers should attempt matching on additional covariates used in neuroimaging, such as age, sex, and a measure of movement (e.g., framewise displacement). Beyond that, using more detailed image quality metrics could be more informative. Processing pipelines, such as MRIqc (Esteban et al., 2017), offer detailed outputs that could provide additional information on quality beyond those offered in the CAT12 toolbox. Ultimately, moving away from subjective visual quality control to automated measures of quality, coupled with PSM, should allow for greater cross-study comparisons, ultimately improving methodological harmonization. To additionally build upon our findings, future research should explore use of PSM in regions of interest or white matter-based analyses as motion is shown to impair morphometric measures across brain regions and tissue types to varying extents (Alexander-Bloch et al., 2016).

In summary, our project demonstrates the use of PSM as a way to increase inclusivity and improve comparability between groups. While impacting datasets differently, this technique improves scan quality imbalance between groups, addressing one of the largest confounds in neuroimaging research.

## CRediT Statement

**Veronica J. Cramm:** Conceptualization, Methodology, Software, Formal Analysis, Investigation, Data Curation, Writing - Original Draft, Writing - Review & Editing, Visualization, Project Administration, Funding Acquisition; **Tyler M. Call:** Conceptualization, Software, Writing - Review & Editing; **John A. E. Anderson:** Conceptualization, Supervision, Funding Acquisition, Project Administration, Resources, Writing - Review & Editing

## Data Acknowledgment

Data used in the preparation of this article were obtained from multiple publicly available neuroimaging initiatives:

- COBRE: Data were downloaded from the COllaborative Informatics and Neuroimaging Suite Data Exchange tool (COINS; http://coins.mrn.org/dx). Data collection was performed at the Mind Research Network and funded by a Center of Biomedical Research Excellence (COBRE) grant 5P20RR021938/P20GM103472 from the NIH to Dr. Vince Calhoun. Please cite: M. Çetin, F. Christensen, C. Abbott, J. Stephen, A. Mayer, J. Cañive, J. Bustillo, G. Pearlson, and V. D. Calhoun, “Thalamus and posterior temporal lobe show greater inter-network connectivity at rest and across sensory paradigms in schizophrenia,” NeuroImage, vol. 97, pp. 117–126, 2014.
- ABIDE: Data used in the preparation of this work were obtained from the Autism Brain Imaging Data Exchange (ABIDE) initiative and downloaded from the COllaborative Informatics and Neuroimaging Suite Data Exchange tool (COINS; http://coins.mrn.org/dx). ABIDE is supported by the Nathan S. Kline Institute for Psychiatric Research, the Child Mind Institute, and other contributing institutions. We gratefully acknowledge the principal investigators and data contributors for making these datasets publicly available. A full list of participating sites and investigators is available at http://fcon_1000.projects.nitrc.org/indi/abide/.
- Neurocognitive Aging Data Release: Data was downloaded from OpenNeuro (https://openneuro.org), dataset ds003592. We thank R. Nathan Spreng and colleagues for publicly releasing the *Neurocognitive Aging* dataset.

This project was supported by an NSERC Discovery Grant (DGECR-2022-00309) and a Canada Research Chair (Tier II, CRC-2020-00174) to JAEA, and an NSERC USRA to VJC (funding reference number 595364).

We have no known conflict of interest to disclose.

## Supplementary Materials

**Table S1.** MRI Acquisition Details for All Three Datasets.

| Dataset | Site | Image Acquisition | Make (model) | Voxel Size (mm) | Flip Angle (deg) | TR (ms) | TE (ms) |
| --- | --- | --- | --- | --- | --- | --- | --- |
| ABIDE | Caltech | 3D MPRAGE | Siemens Magnetom (TrioTim) | 1 x 1 x 1 | 10 | 1590 | 2.73 |
|  | KKI | 3D FFE | Philips (Achieva) | 1 x 1 x 1 | 8 | 8 | 3.7 |
|  | KUL | 3D FFE | Philips (Intera) | 0.98 x 0.98 x 1.2 | 8 | 9.6 | 4.6 |
|  | MPG | 3D MPRAGE | Siemens Magnetom (Verio) | 1 x 1 x 1 | 9 | 1800 | 3.06 |
|  | NYU | 3D MPRAGE | Siemens Magnetom (Allegra) | 1.3 x 1 x 1.3 | 7 | 2530 | 3.25 |
|  | OHSU | 3D MPRAGE | Siemens Magnetom (TrioTim) | 1 x 1 x 1 | 10 | 2300 | 3.58 |
|  | OLIN | 3D MPRAGE | Siemens Magnetom (Allegra) | 1 x 1 x 1 | 8 | 2500 | 2.74 |
|  | Pitt | 3D MPRAGE | Siemens Magnetom (Allegra) | 1.1 x 1.1 x 1.1 | 7 | 2100 | 3.93 |
|  | SBL | 3D FFE | Philips (Intera) | 1 x 1 x 1 | 8 | 9 | 3.5 |
|  | SDSU | 3D SPGR | GE (MR750) | 1 x 1 x 1 | 45 | 11.08 | 4.3 |
|  | SJH | 3D MPRAGE | Siemens Magnetom (Verio) | 1 x 1 x 1 | 8 | 1870 | 2.48 |
|  | Stanford | 3D SPGR | GE (Signa) | 0.86 x 1.5 x 0.86 | 15 | 8.4 | 1.8 |
|  | UCLA | 3D MPRAGE | Siemens Magnetom (TrioTim) | 1 x 1 x 1.2 | 9 | 2300 | 2.84 |
|  | UM | 3D SPGR | GE (MR750) | 1.2 x 1 x 1.2 | 15 | 250 | 1.8 |
|  | USM | 3D MPRAGE | Siemens Magnetom (TrioTim) | 1 x 1 x 1.2 | 9 | 2300 | 2.91 |
|  | Yale | 3D MPRAGE | Siemens Magnetom (TrioTim) | 1 x 1 x 1 | 9 | 1230 | 1.73 |
| **COBRE** | MRN | Multi-Echo<br>MPRAGE | Siemens Magnetom<br>(TrioTim) | 1 x 1 x 1 | 7 | 2530 | 1.64, 3.5,<br>5.36, 7.22,<br>9.08 |
| **Healthy**<br>**Aging** | Cornell | 3D SPGR | GE (MR750) | 1 x 1 x 1 | 7 | 2530 | 3.4 |
|  | York | 3D FLASH | Siemens Magnetom<br>(TrioTim) | 1 x 1 x 1 | 9 | 1900 | 2.52 |

## Footnotes

1 https://openneuro.org/datasets/ds003592/versions/1.0.13 (Healthy Aging) https://fcon_1000.projects.nitrc.org/indi/abide/ (ABIDE) https://fcon_1000.projects.nitrc.org/indi/retro/cobre.html (COBRE)

